# ProteInfer: deep networks for protein functional inference

**DOI:** 10.1101/2021.09.20.461077

**Authors:** Theo Sanderson, Maxwell L. Bileschi, David Belanger, Lucy J. Colwell

**Author notes:** Equal contribution (Theo Sanderson, Maxwell L. Bileschi).

## Abstract

Predicting the function of a protein from its amino acid sequence is a long-standing challenge in bioinformatics. Traditional approaches use sequence alignment to compare a query sequence either to thousands of models of protein families or to large databases of individual protein sequences. Here we instead employ deep convolutional neural networks to directly predict a variety of protein functions – EC numbers and GO terms – directly from an unaligned amino acid sequence. This approach provides precise predictions which complement alignment-based methods, and the computational efficiency of a single neural network permits novel and lightweight software interfaces, which we demonstrate with an in-browser graphical interface for protein function prediction in which all computation is performed on the user’s personal computer with no data uploaded to remote servers. Moreover, these models place full-length amino acid sequences into a generalised functional space, facilitating downstream analysis and interpretation. To read the interactive version of this paper, please visit https://google-research.github.io/proteinfer/

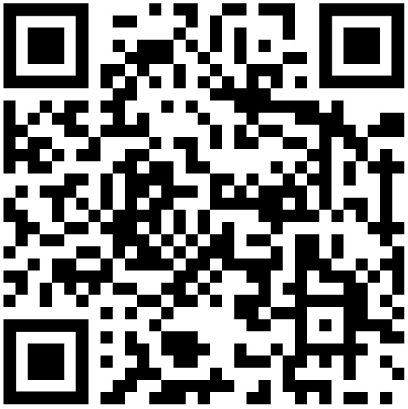

QR code for the interactive version of this preprint at https://google-research.github.io/proteinfer/

## Introduction

Every day, more than a hundred thousand protein sequences are added to global sequence databases (1). However, these entries are of limited use to practitioners unless they are accompanied by functional annotations. While curators diligently extract annotations from the literature, assessing more than 60,000 papers each year (2), the time-consuming nature of this task means that only 0.03% of publicly available protein sequences are manually annotated. The community has a long history of using computational tools to infer protein function directly from amino acid sequence. Starting in the 1980s, methods such as BLAST (3) relied on pairwise sequence comparisons, where a query protein is assumed to have the same function as highly similar sequences that have already been annotated. Signature-based approaches were later introduced, with the PROSITE database (4) cataloguing short amino acid “motifs” found in proteins that share a particular function. Subsequently, a crucial refinement of signature-based approaches was the development of profile hidden Markov models (HMMs) (5, 6). These models collapse an alignment of related protein sequences into a model that provides likelihood scores for new sequences, that describe how well they fit the aligned set. Critically, profile HMMs allow for longer signatures and fuzzier matching and are currently used to update popular databases such as Interpro and Pfam (7, 8). Subsequent refinements have made these techniques more sensitive and computationally efficient (7, 9–12), while their availability as web tools allows practitioners to easily incorporate them into workflows (13–16). These computational modelling approaches have had great impact; however, one third of bacterial proteins still cannot be annotated (even computationally) with a function (17). It is therefore worthwhile to examine how new approaches might complement existing techniques. First, current approaches conduct entirely separate comparisons for each comparator sequence or model, and thus may not fully exploit the features shared across different functional classes. An ideal classification system, for example, might have a modular ATP-binding region detector used in detection of both kinases and ABC transporters (18). Separately modelling these targets in each family, like standard HMM approaches, increases the computational cost and may also be less accurate. In addition, the process of creating many of these signatures is not fully automated and requires considerable curatorial efforts (19, 20), which at present are spread across an array of disparate but overlapping signature databases (8).

Deep neural networks have recently transformed a number of labelling tasks, including image recognition – the early layers in these models build up an understanding of simple features such as edges, and later layers use these features to identify textures, and then entire objects. Edge detecting filters can thus be trained with information from all the labelled examples, and the same filters can be used to detect, for instance, both oranges and lemons (21).

In response, recent work has contributed a number of deep neural network models for protein function classification (22–30). These approaches train a single model to recognise multiple properties, building representations of different protein functions via a series of layers, which allow the same low-level features to be used for different high-level classifications. Of special note is the layer preceding the final layer of the network, which constructs an “embedding” of the entire example in a high-dimensional vector space, and often captures semantic features of the input.

Beyond functional annotation, deep learning has enabled significant advances in protein structure prediction (31–36), predicting the functional effects of mutations (37–40), and protein design (41–47). A key departure from traditional approaches is that researchers have started to incorporate vast amounts of raw, uncurated sequence data into model training, an approach which also shows promise for functional prediction (48).

Of particular relevance to the present work is Bileschi et al. (2019) (49), where it is shown that models with residual layers (50) of dilated convolutions (51) can precisely and efficiently categorise protein domains. Dohan (2021) (52) provides additional accuracy improvements using uncurated data. However, these models cannot infer functional annotations for full-length protein sequences, since they are trained on pre-segmented domains and can only predict a single label. The full-sequence task is of primary importance to biological practitioners.

To address this challenge we employ deep dilated convolutional networks to learn the mapping between full-length protein sequences and functional annotations. The resulting ProteInfer models take amino acid sequences as input and are trained on the well-curated portion of the protein universe annotated by Swiss-Prot (2). We find that: 1) ProteInfer models reproduce curator decisions for a variety of functional properties across sequences distant from the training data, 2) attribution analysis shows that the predictions are driven by relevant regions of each protein sequence, and 3) ProteInfer models create a generalised mapping between sequence space and the space of protein functions, which is useful for tasks other than those for which the models were trained. We provide trained ProteInfer networks that enable other researchers to reproduce the analysis presented and to explore embeddings of their proteins of interest, via both a command line tool^1^, and also via an in-browser JavaScript implementation that demonstrates the computational efficiency of deep-learning approaches.

## Methods

### A neural-network for protein function prediction

In a ProteInfer neural network (Fig. 2), a raw amino acid sequence is first represented numerically as a *one-hot* matrix and then passed through a series of convolutional layers. Each layer takes the representation of the sequence in the previous layer and applies a number of *filters*, which detect patterns of features. We use *residual* layers, in which the output of each layer is added to its input to ease the training of deeper networks (50), and dilated convolutions (51), meaning that successive layers examine larger sub-sequences of the input sequence. After building up an embedding of each position in the sequence, the model collapses these down to a single *n*-dimensional embedding of the sequence using average pooling. Since natural protein sequences can vary in length by at least three orders of magnitude, this pooling is advantageous because it allows our model to accommodate sequences of arbitrary length without imposing restrictive modeling assumptions or computational burdens that scale with sequence length. Finally, a fully-connected layer maps these embeddings to logits for each potential label, which are the input to an element-wise sigmoid layer that outputs per-label probabilities. We select all labels with predicted probability above a given confidence threshold, and varying this threshold yields a tradeoff between precision and recall. To summarize model performance as a single scalar, we compute the F_*max*_ score, the maximum F_1_ score (the geometric mean of precision and recall) across all thresholds (53).

**Fig. 1.**
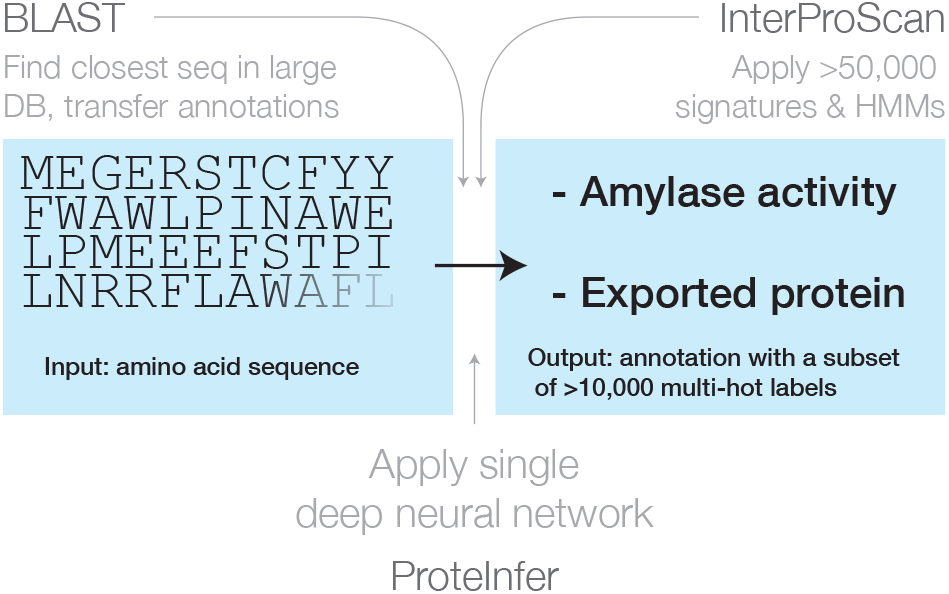
Three approaches for mapping from an amino acid sequence to inferred function: (1) finding similar sequences in a large database of sequences with known annotation (e.g., BLAST), (2) scoring against a large database of statistical models for each family of sequences with known function (e.g. InterProScan), and (3) applying a single deep neural network trained to predict multiple output categories (e.g., this work).

**Fig. 2.**
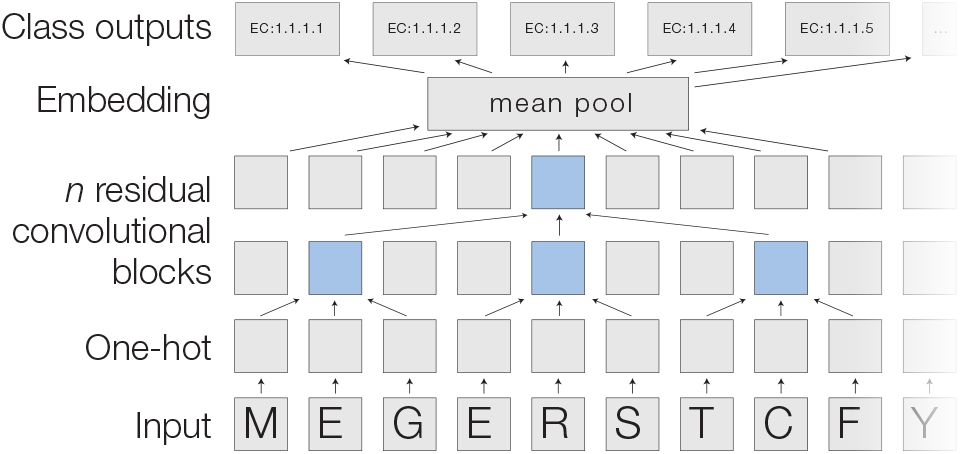
A deep dilated convolutional architecture for protein function prediction. Amino acids are one-hot encoded, then pass through a series of convolutions implemented within residual blocks. Successive filters are increasingly dilated, allowing the top residual layer of the network to build up a representation of high-order protein features. The positional embeddings in this layer are collapsed by mean-pooling to a single embedding of the entire sequence, which is converted into probabilities of each functional classification through a fully connected layer with sigmoidal activations.

Each model was trained for about 60 hours using the Adam optimizer (54) on 8 NVIDIA P100 GPUs with data parallelism (55, 56). We found that using more than one GPU for training improved training time by allowing an increased batch size, but did not have a substantial impact on accuracy compared to training for longer with a smaller learning rate and smaller batch size on one GPU. The models have a small set of hyperparameters, such as the number of layers and the number of filters in each layer, which were tuned using random sampling to maximize F_*max*_ on the random train-test split. Hyperparameter values are available in the Supplement.

### A machine-learning compatible dataset for protein function prediction

The UniProt database is the central global repository for information about proteins. The manually curated portion, Swiss-Prot, is constructed by assessing 60,000 papers each year to harvest 35% of the theoretically curatable information in the literature (2). We focus on Swiss-Prot to ensure that our models learn from human-curated labels, rather than labels generated by a computational annotation pipeline. Each protein in Swiss-Prot goes through a 6-stage process of sequence curation, sequence analysis, literature curation, family-based curation, evidence attribution, and quality assurance. Functional annotation is stored in UniProt largely through *database cross-references*, which link a specific protein with a label from a particular ontology. These cross-references include: Enzyme Commission (EC) numbers, representing the function of an enzyme; Gene Ontology (GO) terms relating to the protein’s molecular function, biological process, or subcellular localisation; protein family information contained in the Pfam (7), SUPFAM(57), PRINTS (58), TIGR (59), PANTHR (60) databases, or the umbrella database InterPro (61); as well as other information including ortholog databases and references on PubMed. Here, we focus on EC and GO labels, though our model training framework can immediately extend to other label sets.

We use two methods to split data into training and evaluation sets. First, a random split of the data allows us to answer the following question: suppose that curators had randomly annotated only 80% of the sequences in Swiss-Prot. How accurately can ProteInfer annotate the remaining 20%? Second, we use UniRef50 (62) clustering to split the data, to model a challenging use-case in which an unseen sequence has low sequence similarity to anything that has been previously annotated. Note there are alternative methods for splitting (48, 63, 64), such as reserving the most recently-annotated proteins for evaluating models. This approach, which is used in CAFA and CASP (63, 64), helps ensure a fair competition because labels for the evaluation data are not available to participants, or the scientific community at large, until after the competition submissions are due. Such a split is not available for EC classification, which is the primary focus of our analyses below. Finally, note that all of the above approaches typically lack reliable annotation for true negatives (65).

To facilitate further development of machine learning methods, we provide TensorFlow (66) TFrecord files for Swiss-Prot ^2^. Each example has three fields: the UniProt accession, the amino acid sequence, and a list of database cross-reference labels. UniProt annotations include only leaf nodes for hierarchical taxononomies such as EC and GO. To allow machine-learning algorithms to model this hierarchy, we added all parental annotations to each leaf node during dataset creation.

## Results

### Prediction of catalysed reactions

We initially trained a model to predict enzymatic catalytic activities from amino acid sequence. This data is recorded as Enzyme Commission (EC) numbers, which describe a hierarchy of catalytic functions. For instance, β amylase enzymes have an EC number of EC:3.2.1.2, which represents the leaf node in the following hierarchy:

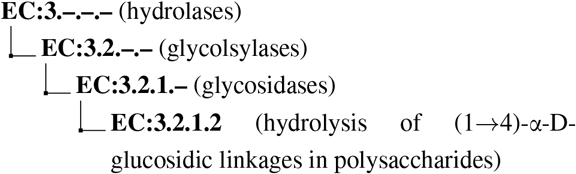

Individual protein sequences can be annotated with zero (non-enzymatic proteins), one (enzymes with a single function) or many (multi-functional enzymes) leaf-level EC numbers. These are drawn from a total of 8,162 catalogued chemical reactions. Our best F_*max*_ was achieved by a model containing 5 residual blocks with 1100 filters each (full details in Supplement). For the dev set, F_*max*_ converged within 500,000 training steps. On the random split, the model achieves F_*max*_ = 0.977 (0.976-0.978) on the held-out test data. At the corresponding confidence threshold, the model correctly predicts 96.7% of true labels, with a false positive rate of 1.4%. Results from the clustered test set are discussed below. Performance was roughly similar across labels at the top of the EC hierarchy, with the highest F_*max*_ score observed for ligases (0.993), and the lowest for oxidoreductases (0.963) (Fig. 3A). For all classes, the precision of the network was higher than the recall at the threshold maximising F_*max*_. Precision and recall can be traded off against each other by adjusting the confidence threshold at which the network outputs a prediction, creating the curves shown in Fig. 3B. BLASTp is arguably a practitioner’s default choice for functional annotation, so we implemented an alignment-based baseline in which BLASTp is used to identify the closest sequence to a query sequence in the train set. Labels are then imputed for the query sequence by transferring those labels that apply to the annotated match from the train set.

**Fig. 3.**
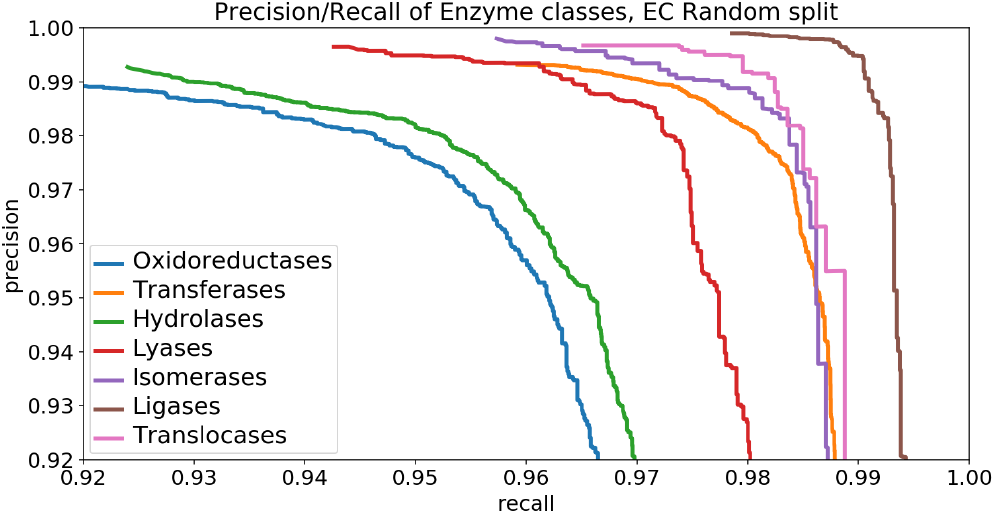
ProteInfer performance for predictions within all 7 top-level enzyme groups.

We produced a precision-recall curve by using the bit score of the closest sequence as a measure of confidence, varying the cutoff above which we retain the imputed labels (63, 67). We also considered an ensemble of neural networks (49), where the average of the ensemble elements’ predicted probabilities is used as a confidence score. (See Fig. S7, Fig. S8.)

We found that BLASTp was able to achieve higher recall values than ProteInfer for lower precision values, while ProteInfer was able to provide greater precision than BLASTp at lower recall values. We wondered whether a combination of ProteInfer and BLASTp could synergize the best properties of both approaches. We found that even the simple ensembling strategy of rescaling the BLAST bit score by the averages of the ensembled CNNs’ predicted probabilities gave a F_*max*_ score (0.991, 95% confidence interval [CI]: 0.990–0.992) that exceeded that of BLAST (0.984, 95% CI: 0.983–0.985) or the ensembled CNN (0.981, 95% CI: 0.980–0.982) alone. On the clustered train-test split based on UniRef50 (see *clustered* in Fig. 3B), we see a performance drop in all methods: this is expected, as remote homology tasks are designed to challenge methods to generalize farther in sequence space. The F_*max*_ score of a single neural network fell to 0.914 (95% CI: 0.913–0.915, precision: 0.959 recall: 0.875), substantially lower than BLAST (0.950, 95% CI: 0.950–0.951), though again an ensemble of both BLAST and ProteInfer outperformed both (0.979, 95% CI: 0.979–0.980). We find that neural network methods learn different information about proteins than alignment-based methods, and a combination of the two further improves remote homology detection.

We also examined the relationship between the number of examples of a label in the training dataset and the performance of the model. In an image recognition task, this is an important consideration since one image of, say, a dog, can be utterly different to another. Large numbers of labels are therefore required to learn filters that are able to predict members of a class. In contrast, for sequence data we found that even for labels that occurred less than five times in the training set, 58% of examples in the test set were correctly recalled, while achieving a precision of 88%, for an F1 of 0.7 (Fig. S9). High levels of performance are maintained with few training examples because of the evolutionary relationship between sequences, which means that one ortholog of a gene may be similar in sequence to another. The simple BLAST implementation described above also performs well, and better than a single neural network, likely again exploiting the fact that many sequence have close neighbours in sequence space with similar functions. We again find that ensembling the BLAST and ProteInfer outputs provides performance exceeding that of either technique used alone.

### Deep models link sequence regions to function

Proteins that use separate domains to carry out more than one enzymatic function are particularly useful in interpreting the behaviour of our model. For example, *S. cerevisiae* fol1 (accession Q4LB35) catalyses three sequential steps of tetrahydrofolate synthesis, using three different protein domains (Fig. 4A). This protein is in our held-out test set, so no information about its labels was directly provided to the model.

**Fig. 4.**
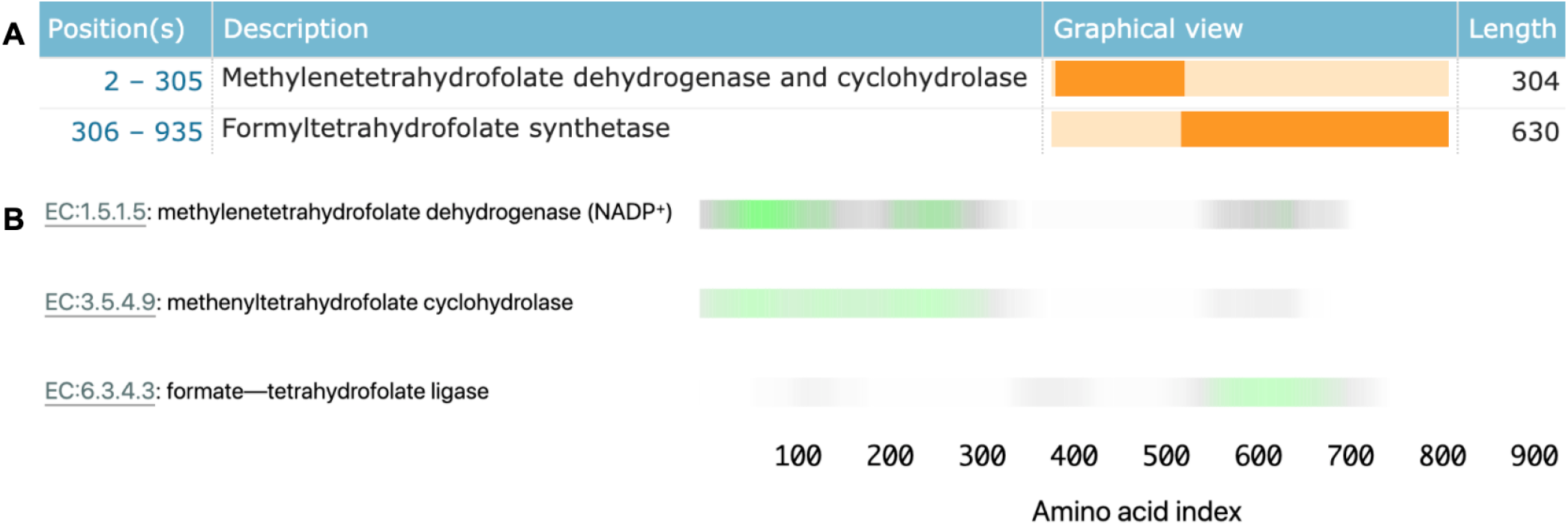
Linking sequence regions to function with class activation mapping for C-1-tetrahydrofolate synthase (accession P11586). A. Ground truth annotation of function on Uniprot (2). B. The three horizontal bars are the sequence region ProteInfer predicts are most involved in each corresponding reaction. This concurs with the known function localization.

To investigate what sequence regions the neural network is using to make its functional predictions, we used class activation mapping (CAM) (68) to identify the sub-sequences responsible for the model predictions. We found that separate regions of sequence cause the prediction of each enzymatic activity, and that these regions correspond to the known functions of these regions (Fig. 4B). This demonstrates that our network identifies relevant elements of a sequence in determining function.

We then assessed the ability of this method to more generally localize function within a sequence, even though the model was not trained with any explicit localization information. We selected all enzymes from Swiss-Prot that had two separate leaf-node EC labels for which our model predicted known EC labels, and these labels were mappable to corresponding Pfam labels. For each of these proteins, we obtained coarse-grained functional localization by using CAM to predict the order of the domains in the sequence and compared to the ground truth Pfam domain ordering(see supplement for details of the method). We found that in 296 of 304 (97%) of the cases, we correctly predicted the ordering, though we note that the set of bifunctional enzymes for which this analysis is applicable is limited in its functional diversity (see supplement). Although we did not find that fine-grained, per-residue functional localization arose from our application of CAM, we found that it reliably provided coarse-grained annotation of domains’ order, as supported by Pfam. This experiment suggests that this is a promising future area for research.

### Neural networks learn a general-purpose embedding space for protein function

Whereas InterProScan compares each sequence against more than 50,000 individual signatures and BLAST compares against an even larger sequence database, ProteInfer uses a single deep model to extract features from sequences that directly predict protein function. One convenient property of this approach is that in the penultimate layer of the network each protein is expressed as a single point in a high-dimensional space. To investigate to what extent this space is useful in examining enzymatic function, we used the ProteInfer EC model trained on the random split to embed each test set protein sequence into a 1100-dimensional vector.

To visualise this space, we selected proteins with a single leaf-level EC number and used UMAP to compress their embeddings into two dimensions (69).

The resulting representation captures the hierarchical nature of EC classification, with the largest clusters in embedding space corresponding to top level EC groupings (Fig. 5A). These clusters in turn are further divided into sub-regions on the basis of subsequent levels of the EC hierarchy (Fig. 5B). Exceptions to this rule generally recapitulate biological properties. For instance, Q8RUD6 is annotated as Arsenate reductase (glutaredoxin) (EC:1.20.4.1) (70) was not placed with other oxidoreductases (EC:1.-.-.-) but rather with Sulfurtransferases (EC:2.8.1.-). Q8RUD6 can however act as a sulfertransferase (71).

**Fig. 5.**
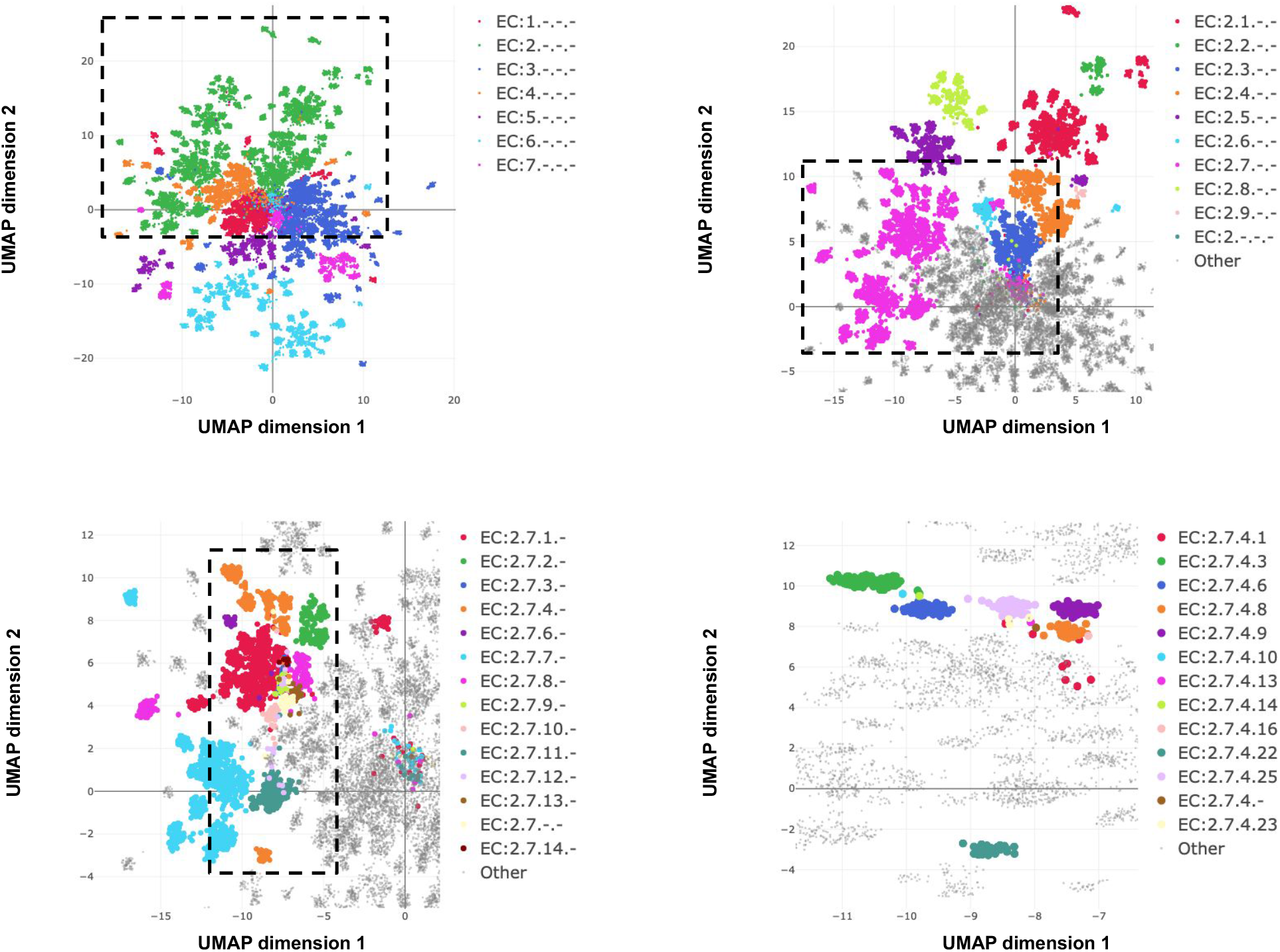
Embedding reflects enzyme functional hierarchy. UMAP projection of embeddings for the subset of test set sequences which have only one leaf-level EC classification. Points are colour-coded at successive levels of the EC hierarchy in each panel. (A) colours denote top level EC groups, (B) colours denote second level EC groups within EC2.*, (C) colours denote third level EC groups within EC:2.7.*, (D) colours depict terminal EC groups within EC:2.7.4.*

Note that the model is directly trained with labels reflecting the EC hierarchy; the structure in Fig. 5 was not discovered automatically from the data. However, we can also ask whether the embedding captures more general protein characteristics, beyond those on which it was directly supervised. To investigate this, we took the subset of proteins in Swiss-Prot that are non-enzymes, and so lack any EC annotations. The network would achieve perfect accuracy on these examples if it e.g. mapped all of them to a single embedding that corresponds to zero predicted probability for every enzymatic label. Do these proteins therefore share the same representation in embedding space? The UMAP projection of these sequences’ embeddings revealed clear structure to the embedding space, which we visualised by highlighting several GO annotations which the network was never supervised on. For example, one region of the embedding space contained ribosomal proteins, while other regions could be identified containing nucleotide binding proteins, or membrane proteins (Fig. 6). To quantitatively measure whether these embeddings capture the function of non-enzyme proteins, we trained a simple random forest classification model that used these embeddings to predict whether a protein was annotated with the *intrinsic component of membrane* GO term. We trained on a small set of non-enzymes containing 518 membrane proteins, and evaluated on the rest of the examples. This simple model achieved a precision of 97% and recall of 60% for an F1 score of 0.74. Model training and data-labelling took around 15 seconds. This demonstrates the power of embeddings to simplify other studies with limited labeled data, as has been observed in recent work (43, 72).

**Fig. 6.**
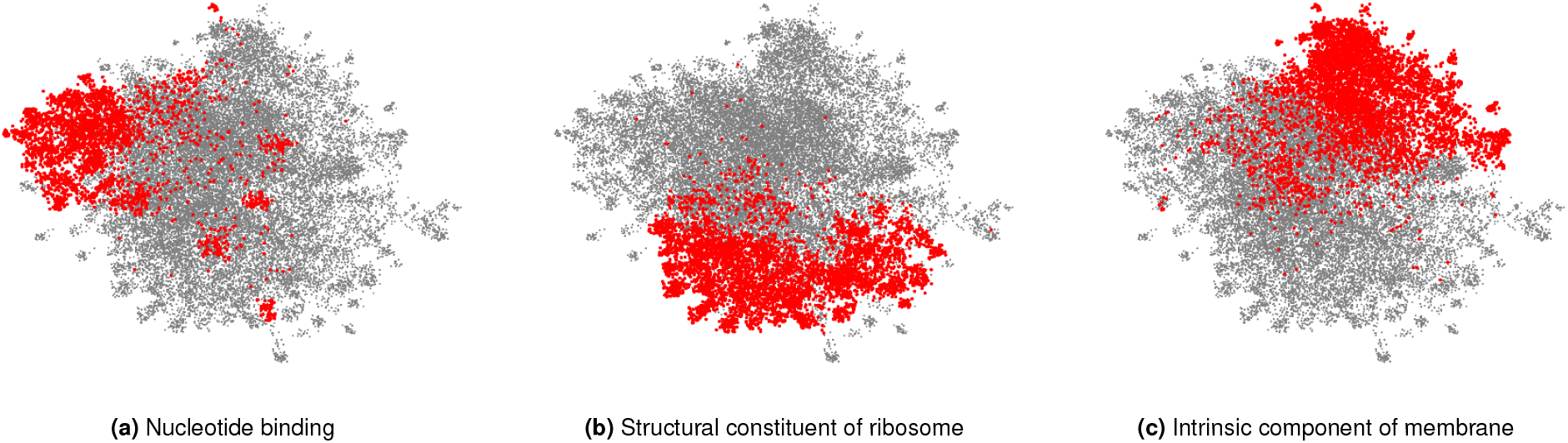
A neural network trained on enzyme function learns general protein properties, beyond enzymatic activity. This figure shows EC-trained ProteInfer embeddings for all non-enzymatic sequences in the test set, projected using UMAP. To illustrate the structure contained in these embeddings we highlight genes based on GO labels (on which this network was never trained).

### Rapid client-side in-browser protein function prediction

Processing speed and ease-of-access are important considerations for the utility of biological software. An algorithm that takes hours or minutes is less useful than one that runs in seconds, both because of its increased computational cost, but also because it allows less immediate interactivity with a researcher. An ideal tool for protein function prediction would require minimal installation and would instantly answer a biologist’s question about protein function, allowing them to immediately act on the basis of this knowledge. Moreover, there may be intellectual property concerns in sending sequence data to remote servers, so a tool that does annotation completely client-side may also be preferable.

There is arguably room for improvement in this regard from classical approaches. For example, the online interface to InterProScan can take 147 seconds to process a 1500 amino acid sequence^3,4^, while running the tool may make the search faster, doing so requires downloading a 9 GB file, with an additional 14 GB for the full set of signatures, which when installed exceeds 51 GB. Meanwhile, conducting a BLAST search against Swiss-Prot takes 34 seconds for a 1500 amino acid sequence ^5^.

An attractive property of deep learning models is that they can be run efficiently, using consumer graphics cards for acceleration. Indeed, recently, a framework has been developed to allow models developed in TensorFlow to be run locally using simply a user’s browser (73), but to our knowledge this has never been deployed to investigate biological sequence data. We therefore built a tool to allow near-instantaneous prediction of protein functional properties in the browser. When the user loads the tool, lightweight EC (5MB) and GO model (7MB) prediction models are downloaded and all predictions are then performed locally, with query sequences never leaving the user’s computer. Inference in the browser for a 1500 amino-acid sequence takes < 1.5 seconds for both models (Sup. Note G).

### Comparison to experimental data

To assess our model’s performance using an additional experimentally-validated source of ground truth, we focused our attention on a large set of bacterial genes for which functions have recently been identified in a high-throughput experimental genetic study (17). In particular, this study listed newly identified EC numbers for 171 proteins, representing cases when there was previously either misannotation or inconsistent annotation in the SEED or KEGG databases.

Therefore, this set of genes may be enriched for proteins whose functions are difficult to assess computationally. We examined how well our network was able to make predictions for this experimental dataset at each level of the EC hierarchy (Fig. 7), using as a decision threshold the value that optimised F1 identified during tuning. The network had high accuracy for identification of broad enzyme class, with 90% accuracy at the top level of the EC hierarchy. To compute accuracy, we examined the subset of these 171 proteins for which there was a single enzymatic annotation from (17), giving us predictions for 119 enzymes. At the second level of the hierarchy, accuracy was 90% and the network declined to make a prediction for 12% of classes. Even at the third level, accuracy was 86% with the network making a prediction in 77% of cases. At the finest level of classification, the proportion of examples for which a prediction was made fell to 28%, with 42% of these predictions correct.

**Fig. 7.**
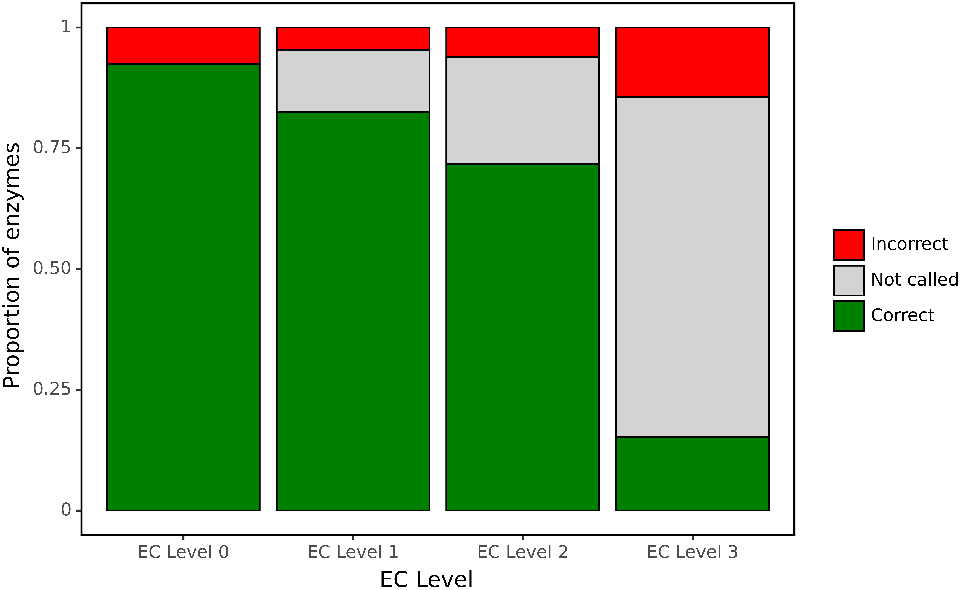
ProteInfer predictions for a set of genes recently experimentally reannotated by high-throughput phenotyping. ProteInfer makes confident and largely accurate predictions at the earliest levels of the EC hierarchy. Accuracy falls at the finest levels of classification (for this set of challenging genes) but fortunately the network declines to make a prediction in most cases, with every label failing to meet the threshold for positive classification.

As an example, the *Sinorhizobium meliloti* protein Q92SI0 is annotated in UniProt as a Inosine-uridine nucleoside N-ribohydrolase (EC 3.2.2.1). Analysing the gene with InterProScan (74) also gives this prediction, but our model instead predicts it to be a uridine nucleosidase (EC 3.2.2.3), and this was indeed the result found in this experimental work. Similarly, *Pseudomonas fluorescens* A0A166Q345 was correctly classified by our model as a D-galacturonate dehydrogenase (EC 1.1.1.203) as opposed to a misannotation on UniProt and with InterProScan.

It was notable that for many of these proteins, the network declined to make a prediction at the finest level of the EC hierarchy. This suggests that by training on this hierarchical data, the network is able to appropriately make broad or narrow classification decisions. This is similar to the procedure employed with manual annotation: when annotators are confident of the general class of reaction that an enzyme catalyses but not its specific substrate, they may leave the third or fourth position of the EC number blank (e.g. EC:1.1.-.-). Due to training on hierarchical data, our network is able to reproduce these effects by being more confident (with higher accuracy) at earlier levels of classification.

### A model predicting the entire gene ontology

Given the high accuracy that our deep learning model was able to achieve on the more than five thousand enzymatic labels in Swiss-Prot, we asked whether our networks could learn to predict protein properties using an even larger vocabulary of labels, using a similar test-train setup. Gene Ontology (75–77) (GO) terms describe important protein functional properties, with 28,079 such terms in Swiss-Prot that cover the molecular functions of proteins (e.g. DNA-binding, amylase activity), the biological processes they are involved in (e.g. DNA replication, meiosis), and the cellular components to which they localise (e.g. mitochondrion, cytosol). These terms are arranged in a complex directed acyclic graph, with some nodes having as many as 12 ancestral nodes.

We note that there has been extensive work in GO label prediction evaluated on a temporally-split dataset (constructing a test set with the most recently experimentally annotated proteins), e.g., (63), and stress that our comparison is based on the random and clustered splits of Swiss-Prot described above. This approach to splitting the data into train and test has advantages and disadvantages as compared to a temporal split, which depend on the desired application for the method being evaluated.

We trained a single model to predict presence or absence for each of these terms and found that our network was able to achieve a precision of 0.918 and a recall of 0.854 for an F1 score of 0.885 (95% CI: 0.882–0.887).

An ensemble of multiple CNN elements was again able to achieve a slightly better result with an F1 score of 0.899 (95% CI: 0.897–0.901), which was exceeded by a simple transfer of the BLAST top pick at 0.902 (95% CI: 0.900–0.904), with an ensemble of both producing the best result of 0.908 (95% CI: 0.906–0.911).

To benchmark against a common signature-based methodology, we used InterProScan to assign protein family signatures to each test sequence. We chose InterProScan for its coverage of labels as well as its use of multiple profile-based annotation methods, including HMMER and PROSITE, mentioned above. We note that while InterProScan predicts GO labels directly, it does not do so for EC labels, which is why we did not use InterProScan to benchmark our work on predicting EC labels. We found that InterProScan gave good precision, but within this UniProt data had lower recall, giving it a precision of 0.937 and recall of 0.543 for an F1 score of 0.688. ProteInfer’s recall at a precision of .937 is substantially higher (0.835) than InterProScan at assigning GO labels.

There are multiple caveats to these comparisons. One challenge is that the completeness of Swiss-Prot’s GO term annotations varies (78). As an extreme example, *Pan paniscus* (Pygmy Chimpanzee) and *Pan troglodytes* (Chimpanzee) have an identical Apolipoprotein A-II protein,^6^, where the first protein has 24 GO annotations, while the latter has 143 GO annotations.^7^ One way this is reflected in the performance of the models is that some BLAST matches that have extremely large bit-scores are not annotated identically, and thus reduce the precision of the BLAST model. It is also important to note that our model has the advantage of being specifically trained on the UniProt labelling schema upon which it is being evaluated. InterPro works quite differently, with GO terms being assigned to families, and so inconsistencies in terms of how these are assigned can explain reduced performance – for instance InterPro families simply do not feature all of the GO terms found in UniProt. Thus these results should be seen as specific to the task of reproducing the curated results in UniProt.

## Discussion

We have shown that neural networks trained and evaluated on high-quality Swiss-Prot data accurately predict functional properties of proteins using only their raw, un-aligned amino acid sequences. Further, our models make links between the regions of a protein and the function that they confer, produce predictions that agree with experimental characterisations, and place proteins into an embedding space that captures additional properties beyond those on which the models were directly trained. We have provided a convenient browser-based tool, where all computation runs locally on the user’s computer. To support follow-up research, we have also released our datasets, code for model training and evaluation, and a command-line version of the tool.

Using Swiss-Prot to benchmark our tool against traditional alignment-based methods has distinct advantages and disadvantages. It is desirable because the data has been carefully curated by experts, and thus it contains minimal false-positives. On the other hand, many entries come from experts applying existing computational methods, including BLAST and HMM-based approaches, to identify protein function. Therefore, the data may be enriched for sequences with functions that are easily ascribable using these techniqueswhich could limit the ability to estimate the added value of using an alternative alignment-free tool. An idealised dataset would involved training only on those sequences that have themselves been experimentally characterized, but at present too little data exists than would be needed for a fully supervised deep-learning approach. Semi-supervised approaches that combine a smaller number of high quality experimental labels with the vast set of amino acid sequences in TrEMBL may be a productive way forward.

Further, our work characterizes proteins by assigning labels from a fixed, pre-defined set, but there are many proteins with functions that are not covered by this set. These categories of functions may not even be known to the scientific community yet. There is a large body of alternative work that identifies groups of related sequences (e.g. (79)), where a novel function could be discovered, for example, using follow-up experiments.

Finally, despite the successes of deep learning in many application domains, a number of troublesome behaviours have also been identified. For example, probabilities output by deep models are often over-confident, rather than well-calibrated (80), and networks perform poorly on out-of-distribution data without being aware that they are outside their own range of expertise (81). Though these issues still need to be addressed and better understood by both the machine learning and bioinformatics communities, deep learning continues to make advances in a wide range of areas relating to the understanding protein function. We thus believe deep learning will have a central place in the future of this field.

Our code, data, and notebooks reproducing the analyses shown in this work are available online at https://github.com/google-research/proteinfer and https://console.cloud.google.com/storage/browser/brain-genomics-public/research/proteins/proteinfer/datasets/.

## ACKNOWLEDGEMENTS

We would like to thank Babak Alipanahi, Jamie Smith, Eli Bixby, Drew Bryant, Shanqing Cai, Cory McLean and Abhinay Ramparasad. The static version of this manuscript uses a template made by Ricardo Henriques.

TS receives funding from the Wellcome Trust through a Sir Henry Wellcome Post-doctoral Fellowship (210918/Z/18/Z). This work was also supported by the Francis Crick Institute which receives its core funding from Cancer Research UK (FC001043), the UK Medical Research Council (FC001043), and the Wellcome Trust (FC001043). This research was funded in whole, or in part, by the Wellcome Trust [FC001043]. For the purpose of Open Access, the authors have applied a CC BY public copyright licence to any Author Accepted Manuscript version arising from this submission.

## AUTHOR CONTRIBUTIONS

TS was responsible for conception, data ingestion, code, analysis, visualisation, UI, paper writing. MLB was responsible for visualization, analysis, code, CLI, open sourcing, testing, paper writing. DB was responsible for paper writing, analysis of results, code reviews. LJC was responsible for problem conception and framing, analysis of results, paper writing.

## Footnotes

1 https://github.com/google-research/proteinfer

2 https://console.cloud.google.com/storage/browser/brain-genomics-public/research/proteins/proteinfer/datasets/

3 Protein Q77Z83.

4 It should be noted that the individual databases that make up InterProScan may return matches faster, with the online interface to Pfam taking 14–20 seconds for a 1500 amino acid sequence.

5 A target database with decreased redundancy could be built to reduce this search time, and other optimizations of BLAST have been developed.

6 Accessions P0DM95 and Q8MIQ5.

7 This count is done using not only the set of all labels that appear in Swiss-Prot, but also any parents of those labels.

## Notes

https://google-research.github.io/proteinfer/

https://github.com/google-research/proteinfer

### Competing Interest Statement

The authors have declared no competing interest.

