## Supplement for "ProteInfer: deep networks for protein functional inference"

### A. Implementation details

***Label inheritance.*** Some annotations, such as GO terms and EC numbers, are structured as directed acyclic graphics (which take the form of simple trees for EC numbers). Typically in such cases the annotation provided on UniProt is the most-specific that is known. For example if a protein is known to exhibit *sequence specific DNA binding* (GO:0043565). It will not separately be annotated with the ancestral term *DNA binding* (GO:0003677), this is simply assumed from the ontology. Using such an annotations directly, however, is likely to be problematic in a deep learning setting. Failing to annotate an example with the parental term demands that the model predict that the example is negative for this term, which is not the effect we want. To address this our datasets include labels for all ancestors of applied labels for EC, GO and InterPro datasets. In the case of GO we restrict these to *is_a* relationships.

***Class activation mapping.*** Many proteins have multiple functional properties. For example we analyse the case of the bifunctional dhfr/ts of *T. gondii*. Such bifunctional enzymes are often not unique – it is functionally advantageous for these enzymes to be fused, which facilitates channeling of substrate between their active sites. Since there are a number of such examples within Swiss-Prot, the mere existence of a TS domain in a protein is (mild) evidence for possible DHFR function. To increase the interpretability of the network we subtract the activations for other predicted classes from those of the class of interest during class activation mapping.

#### Model architecture

To create an architecture capable of receiving a wide range of input sequences, with computational requirements determined for each inference by the length of the individual input sequence, we employed a dilated convolutional approach (51). Computation for both training and prediction in such a model can be parallelized across the length of the sequence. By training on full length proteins, in a multi-label training setting, we aimed to build networks that could extract functional information from raw amino acid sequences. One helpful feature of this architecture is its flexibility with regards to sequence length. Natural protein sequences can vary in length by at least three orders of magnitude, but some architectures have computational requirements that scale with the maximum sequence they are capable of receiving as input, rather than the sequence being currently examined. These fixed-length approaches reduce efficiency as well as place a hard limit on the length of sequences that can be examined.

### B. Input data statistics

We use Swiss-Prot version 2019_01 in our analysis, which gives us 559077 proteins, or 548264 after filtering for only 20 standard amino acids and filtering fragments. Because different protein functions have differing prevalence, we note the

| Fold | Number of sequences |
|---|---:|
| train | 438522 |
| dev | 55453 |
| test | 54289 |
| all together | 548264 |

**Table S1.** In our random split of the training data, we allocate about 80% to the training fold, 10% to the development fold, and 10% to the test fold.

| Fold | Number of sequences |
|---|---:|
| train | 182965 |
| dev | 180309 |
| test | 183475 |
| all together | 546749 |

**Table S2.** In our clustered split of the training data, we use UniRef50, and allocate approximately equal numbers of sequences to each fold.

number of proteins that have a given function for Pfam, EC, and GO labels, as well as noting the number of labels per protein.

When assigning examples to folds in our clustered dataset, we note that there are test examples that have labels that are never

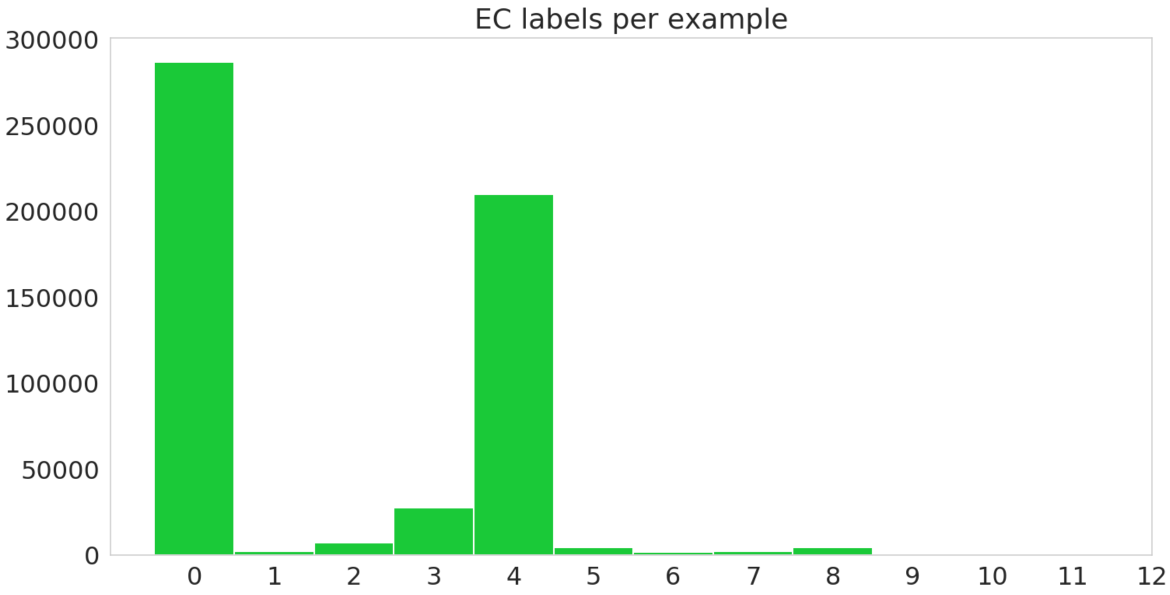


**Fig. S1.** Histogram of number of labels per sequence, including hierarchical labels, on the random dataset.

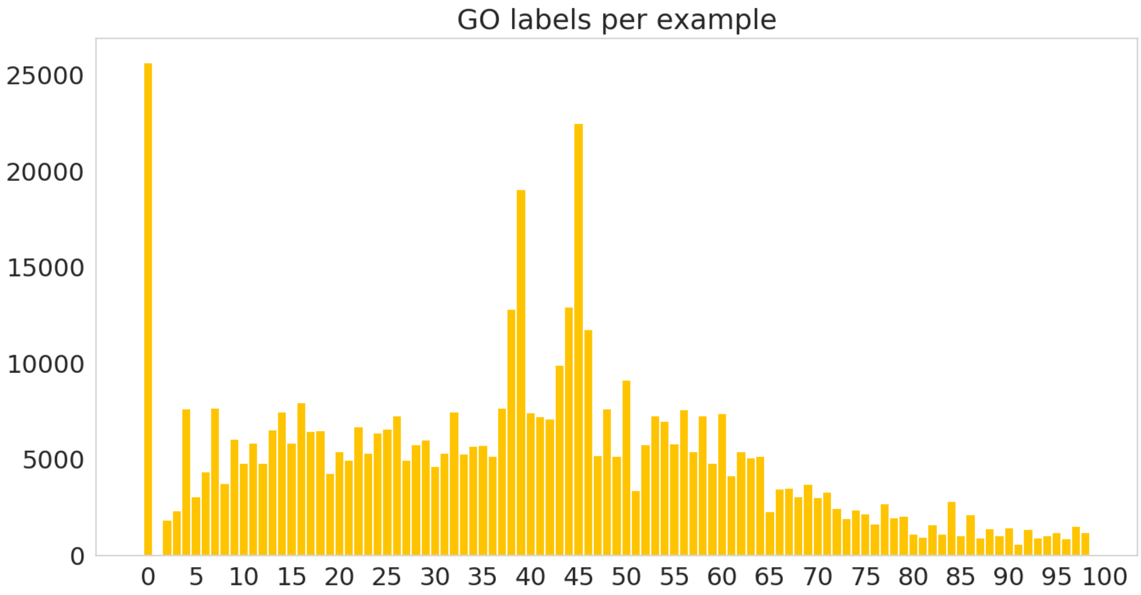


**Fig. S2.** Histogram of number of labels per sequence, including hierarchical labels, on the random dataset.

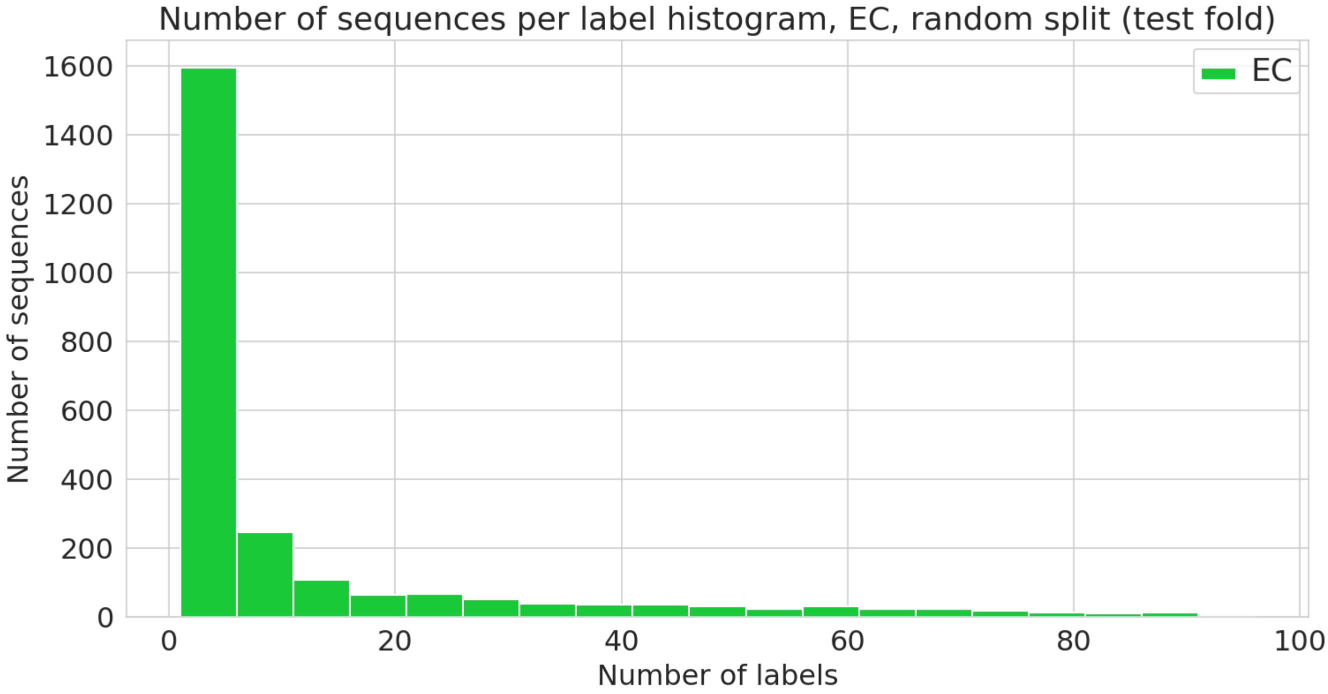


**Fig. S3.** Number of sequences annotated with a given functional label. (EC class) in the random dataset.

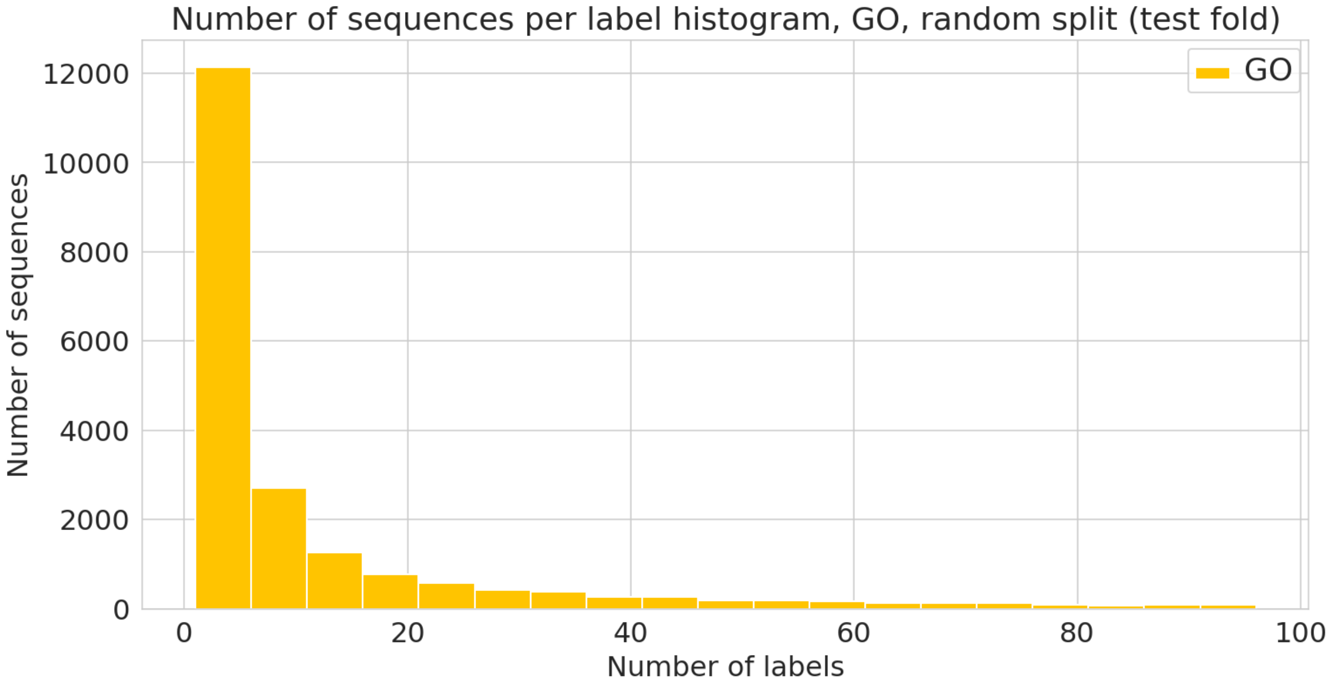


**Fig. S4.** Number of sequences annotated with a given functional label. (GO label) in the random dataset.

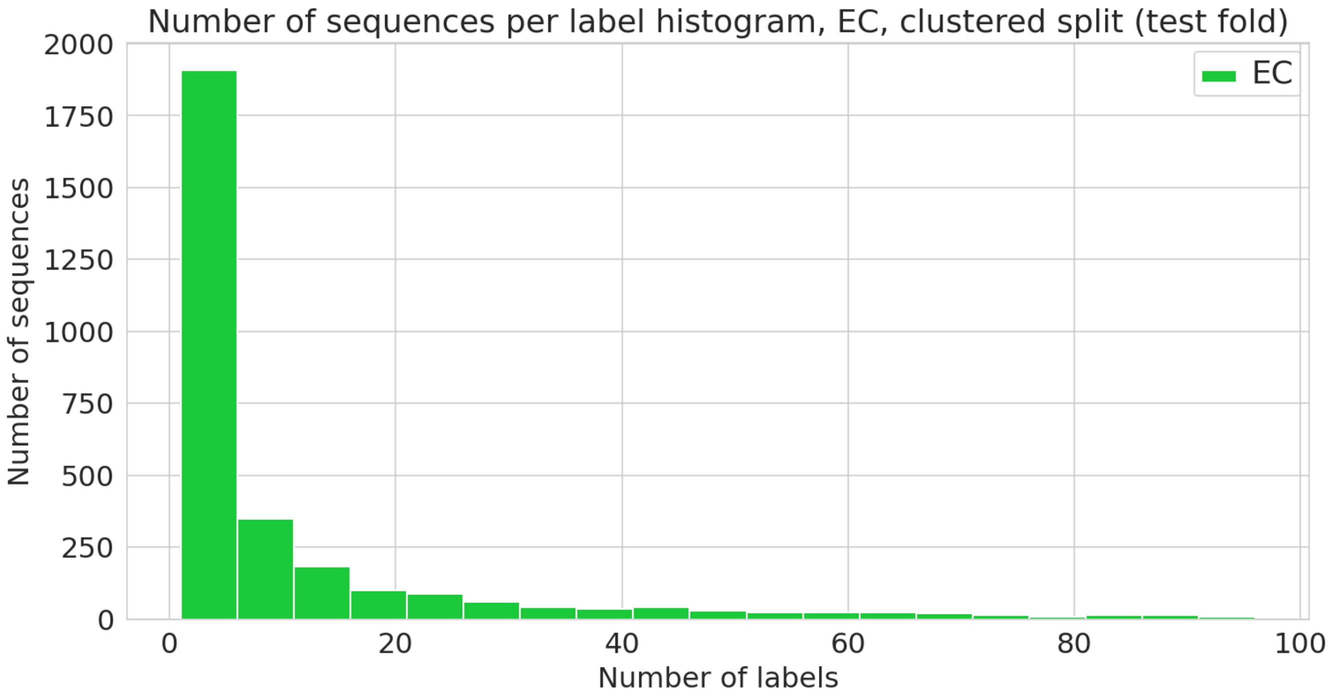


**Fig. S5.** Number of sequences annotated with a given functional label. (EC class) in the clustered dataset.

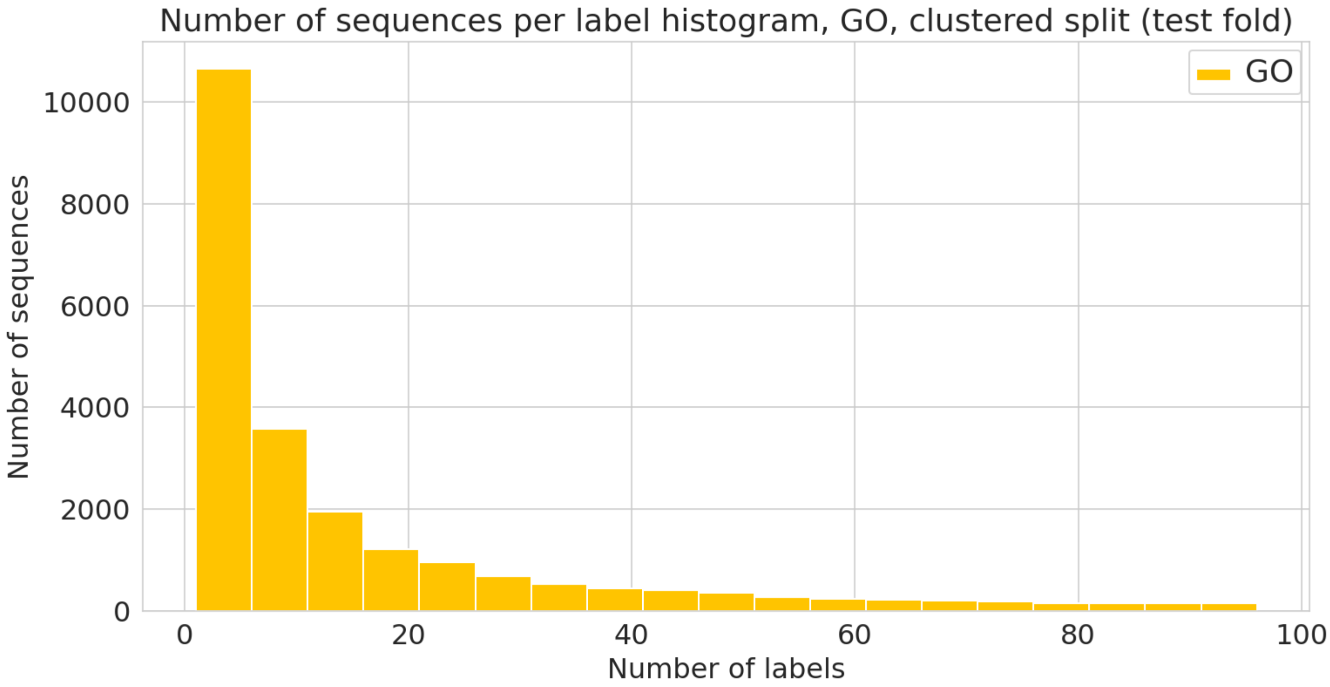


**Fig. S6.** Number of sequences annotated with a given functional label. (GO label) in the clustered dataset.

seen in the training data. We report these cases below as "Impossible" test example-label pairs.

| Type | Number |
|---|---|
| Train labels | 3411 |
| Test labels | 3414 |
| Impossible test labels | 1043 |
| Train example-label pairs | 348105 |
| Test example-label pairs | 348755 |
| Impossible test example-label pairs | 3415 |

**Table S3.** Clustered dataset statistics for EC labels.

| Type | Number |
|---|---|
| Train labels | 26538 |
| Test labels | 26666 |
| Impossible test labels | 3739 |
| Train example-label pairs | 8338584 |
| Test example-label pairs | 8424299 |
| Impossible test example-label pairs | 11137 |

**Table S4.** Clustered dataset statistics for GO labels.

### C. Precision/Recall curves

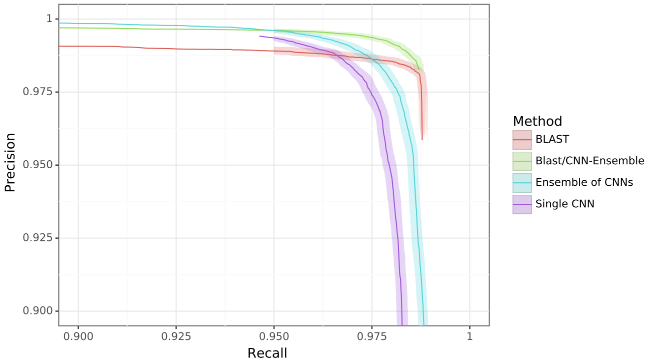


**(a)** EC number prediction: random split

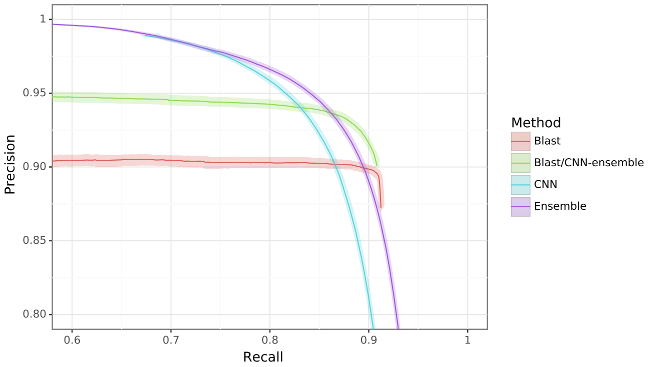


**(b)** GO term prediction: random split

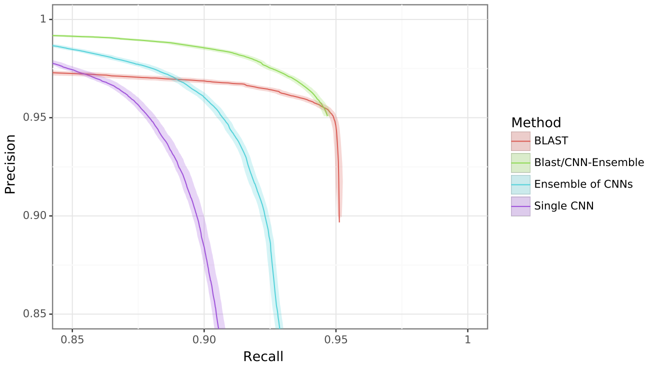


**(c)** EC number prediction: clustered split

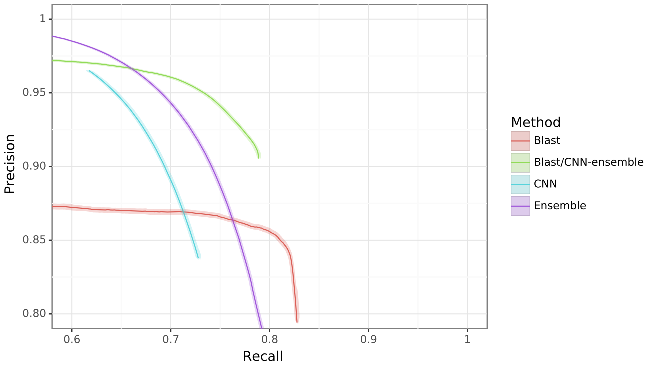


**(d)** GO term prediction: clustered split

**Fig. S7.** Bootstrapped precision-recall curves for EC number prediction and gene ontology term prediction for random and clustered splits for four methods: BLAST top pick, single ProteInfer CNN, ensembled ProteInfer CNNs, and ensembled ProteInfer CNNs scaled by BLAST score.

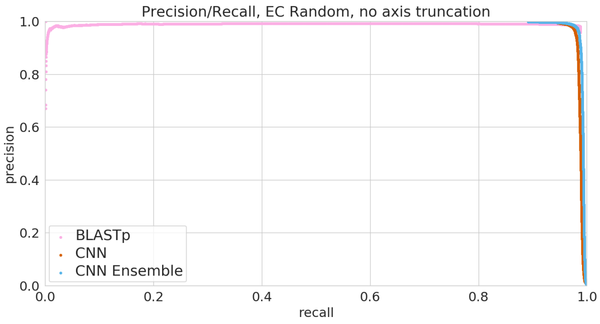


**(a)** EC number prediction: random split

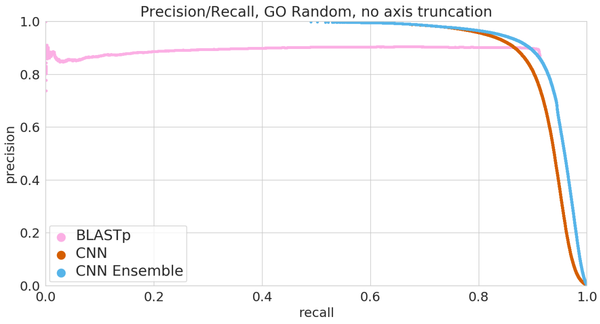


**(b)** GO term prediction: random split

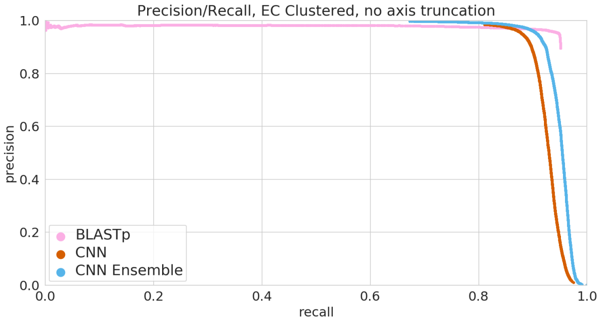


**(c)** EC number prediction: clustered split

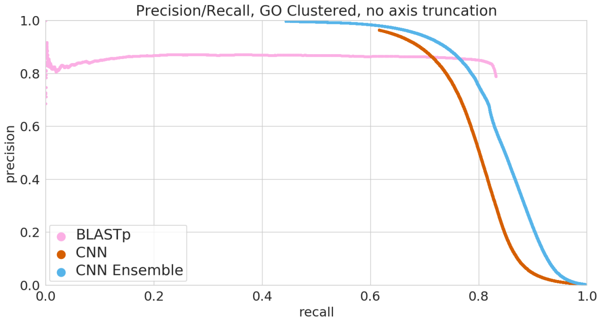


**(d)** GO term prediction: clustered split

**Fig. S8.** Full precision-recall curves for EC number prediction and gene ontology term prediction for random and clustered splits for four methods: BLAST top pick, single ProteInfer CNN, ensembled ProteInfer CNNs

### D. Stratified performance

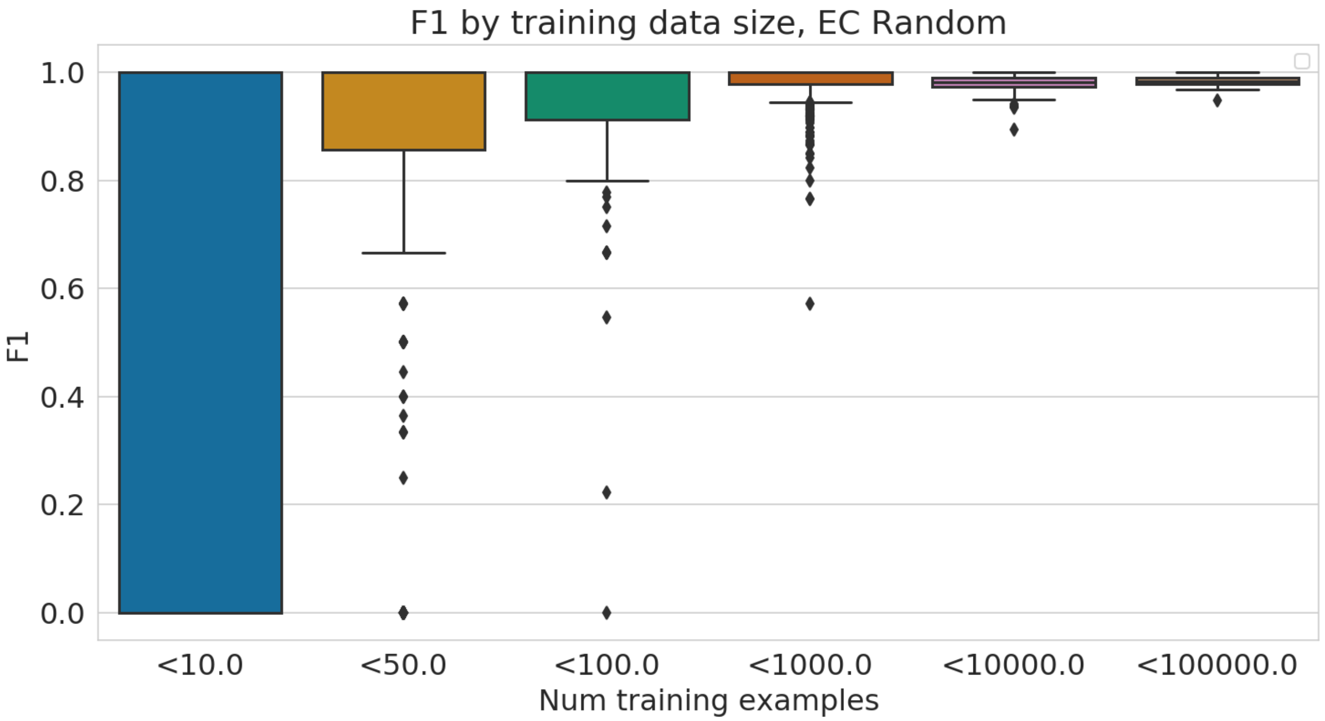


**Fig. S9.** Performance of EC model stratified by number of training examples available for each test example.

### E. Hyperparameters

We tuned over batch size, dilation rate, filters, first dilated layer, kernel size, learning rate, number of layers, mean vs max pooling, and Adam $\beta_1$, $\beta_2$ and $\epsilon$ (54) over a number of studies to determine the set of parameters that optimized $\texttt{F}_{max}$. We found, as in (49), that the network was relatively unresponsive to slight changes in hyperparameters, and found that many of the hyperparameters that performed well in (49) also performed well for this task. We chose to keep identical hyperparameters for the EC and GO tasks across both the random and clustered splits for simplicity, and we note that parameters with good performance on the random task performed respectively well on the clustered split.

| | CNN |
|---|---|
| concurrent batches (data parallelism) | 8 |
| batch size | 40 (per each GPU)<br>Dynamic based on sequence length |
| dilation rate | 3 |
| filters | 1100 |
| first dilated layer | 2 |
| gradient clip | 1 |
| kernel size | 9 |
| learning rate | 1.5E-3 |
| learning rate decay rate | 0.997 |
| learning rate decay steps | 1000 |
| learning rate warmup steps | 3000 |
| Adam $\beta_1$ | .9 |
| Adam $\beta_2$ | .999 |
| Adam $\epsilon$ | 1E-8 |
| number of ResNet layers | 5 |
| pooling | mean |
| ResNet bottleneck factor | 0.5 |
| train steps | 500000 |

**Table S5.** Hyperparameters used in convolutional neural networks. We note that hyperparameters for single-GPU training are available in `github.com/google-research/proteinfer/blob/master/hparams_sets.py`.

### F. Predicting coarse-grained functional localization with CAM

The goal of this experimental methodology is to measure whether or not we correctly order the localization of function in bifunctional enzymes. As such, first we have to identify a set of candidates for experimentation.

***F.1. Candidate set construction.*** We note that no functional localization information was available to our models during training, so we can consider not just the dev and test sets, but instead the entirety of Swiss-Prot for our experimentation. As such, we take all examples from Swiss-Prot that have an EC label, and convert these labels to Pfam labels using a set of 1515 EC-Pfam manually curated label correlations from InterPro (82), omitting unmapped labels. We then take the set of 3046 proteins where exactly two of their ground-truth labels map to corresponding Pfam labels. In our Swiss-Prot random test-train split test set, on bifunctional enzymes, we get $0.995$ precision and $0.948$ recall at a threshold of $0.5$, so we believe this set is a reasonable test set for ordering analysis.

We then predict EC labels for these proteins with one of our trained convolutional neural network classifier, considering only the most specific labels in the hierarchy. Then, we map these predicted EC labels to Pfam labels using the InterPro mapping again, and retain only the proteins on which we predict exactly two labels above a threshold of $0.5$, and are left with 2679 proteins. In 2669 out of 2679 proteins, our predictions are identical to the Pfam-mapped ground truth labels.

We take these 2669 that have two true and predicted *Pfam* labels, and look at their current Pfam labels annotated in Swiss-Prot. Of these 2669 proteins, 304 of them contain both of the mapped labels. We note that this difference between 2669 and 304 is likely due in part to Pfam being conservative in calling family members, potential agreements at the Pfam clan vs family level, as well as database version skew issues.

***F.2. Computation of domain ordering.*** On these 304 proteins we have the same predicted-EC-to-Pfam labels and the same true-Pfam labels. For each of these proteins, we can get an ordering of their two enzymatic domains from Pfam, giving us a *true* ordering. It is now our task to produce a predicted ordering.

We use class activation mapping (CAM) to compute a confidence for each class at each residue for every protein in this set of 304. We then filter this large matrix of values and only consider the families for which our classifier predicted membership, giving us a matrix of shape sequence-length by predicted-classes (which is two in this case). For each class, we take the CAM output and compute a center of mass. Then we order the two classes based on where their center of mass lies.

| First domain | Second Domain | Number ordered correctly | Number times seen | Percent Correct |
|---|---|---|---|---|
| EC:2.7.7.60 | EC:4.6.1.12 | 94 | 94 | 100 |
| EC:4.1.99.12 | EC:3.5.4.25 | 83 | 83 | 100 |
| EC:3.5.4.19 | EC:3.6.1.31 | 59 | 59 | 100 |
| EC:1.8.4.11 | EC:1.8.4.12 | 20 | 20 | 100 |
| EC:4.1.1.48 | EC:5.3.1.24 | 18 | 18 | 100 |
| EC:5.4.99.5 | EC:4.2.1.51 | 12 | 12 | 100 |
| EC:5.4.99.5 | EC:1.3.1.12 | 4 | 4 | 100 |
| EC:4.2.1.10 | EC:1.1.1.25 | 3 | 3 | 100 |
| EC:2.7.7.61 | EC:2.4.2.52 | 0 | 3 | 0 |
| EC:2.7.1.71 | EC:4.2.3.4 | 0 | 2 | 0 |
| EC:1.1.1.25 | EC:4.2.1.10 | 0 | 1 | 0 |
| EC:2.7.2.3 | EC:5.3.1.1 | 1 | 1 | 100 |
| EC:4.1.1.97 | EC:1.7.3.3 | 1 | 1 | 100 |
| EC:4.1.3.1 | EC:2.3.3.9 | 1 | 1 | 100 |
| EC:5.1.99.6 | EC:1.4.3.5 | 0 | 1 | 0 |
| EC:1.8.4.12 | EC:1.8.4.11 | 0 | 1 | 0 |

**Table S6.** Domain architecture diversity in bifunctional enzymes. In Swiss-Prot, there are 16 candidate domain architectures available for our EC functional localization experiment. Among these all domain architectures with more than 3 instances in Swiss-Prot (7 of them) are 100% correctly ordered by our CAM method.

### G. Timing ProteInfer browser models

In the javascript console, we run the following:

```
var seq = "MEPAKPSGNNMGSNDERMQDYRPDPMMEESIKEILEESLMCDTSFDDLIIPGLESFGLIIPESSNNIESNNVEEGSDGE" +
"LKTLAEQKCKQGNDNDVIQSAMKLSGLYCDADITHTQPLSDNTHQDPIYSQESRIFTKTIQDPRIVAQTHRQCTSSASNL" +
"QSNESGSTQVRFASELPNQLLQPMYTSHNQNANLQNNFTSLPYQPYHDPYRDIESSYRESRNTNRGYDYNFRHHPYRPRG" +
"GNGKYNYYNPNSKYQQPYKRCFTRTYNRRGRGHRSYDCSDRSADLPYEHYTYPNYEQQNPDPRMNNYKDFTQLTNKFNFE" +
"SYDYSMAFSTDSTHVQSDNYNHPTKAQTIPETTKTKKHEATKDNETSTENQVLTPDVISLSYRPSSYKMDIIKKIYDTDV" +
"IPLPKEALTANGSNRDVDIQKYKKAHIRCRSVQKKKERSSQTNKHDENHASSRSDLKERKSNENEDKAVTKARDFSKLNP" +
"LLSPLPLTPEPAIDFADHTDKFYSTPEFNQIQQNLHRSKTSLQDTVPISKHTPRAPTKDNSYKKHHDSKDNYPKMKHSPG" +
"RTTSKKNTTNSNGHQNFKEVSVKNVSGKATSTSPKSKTHHYSSSSDEEGQYKSPVKTIIQSPSPYCKLKNPSIMDKNSAK" +
"NHTASADKNLTDNSPIRSNLNPTAFNKSNSNKSITDSTSNSDECTDRKPNCNSTKNESKDPNRTCGKNSDKHLSKSCTMA" +
"SKRAPSRASSRASSRDSSRASSRASSRASSRDSSRASSRASSRDSSRASSRASSRASSKASSRASSRASSRASSRDSSRA" +
"SSKASSRASSRDSSRASSRASSRASSKASSRASSRASSRASSRDSSRASSKASSRASSRDSSRASSRDSSRDSSRASSRA" +
"SSRDSSRASSKASRKASSRASSRASSRASSKASGKASSEASSRASSRNSSRASSRASSRASSRDSSRASSRASSRDSSRA" +
"SSKASRKASSRASSRASSRELRQIYCDSNKRQTPPHDTSINTKFEISEIKFRCGEDLNFYKNTAARLQCFNHNDQFYNPR" +
"FRPHIRTNRKKSESTNDTDSESSMSRCKSHCRNSPDSLTVVRRKKHKSGSSSISSSIEENCRSNSHIVTGKEKFTPFYYQ" +
"SSRTRSSSSSSSSSASLSCSKSTLKTCRKTQYKDNKQIKSKSDSKHKTTNMSSDYESNRHADVFKNSPEAGEKFPLHNSS" +
"PFNTHEQSNHSENAIDEEQKKAPNITTSHLQGKQNVRLHNTKKCKKKRPRDDDSDSSIKNFCKKRISGAQKTESEVSEPD" +
"DLCYRDYVRLKERKVSEKFKIHRGRVATKDFQKLFRNTMRAFEYKQIPKKPCNEKNLKEAVYDICCNGLSNNAAIIMYFT" +
"RSKKVAQIIKIMQKELMIRPNITVSEAFKMNHAPPKYYDKDEIKRFIQLQKQGPQELWDKFENNTTHDLFTRHSDVKTMI" +
"IYAATPIDFVGAVKTCNKYAKDNPKEIVLRVCSIIDGDNPISIYNPISKEFKSKFSTLSKC"
var t0 = performance.now()
await Promise.all([performECInference(seq), performGOInference(seq)])
var t1 = performance.now()
console.log("Call took " + (t1 - t0) + " milliseconds.")
```
